# Quantifying the potential for future gene therapy to lower lifetime risk of polygenic late-onset diseases

**DOI:** 10.1101/390773

**Authors:** Roman Teo Oliynyk

## Abstract

**Background:** Within the next few decades, gene therapy techniques and genetic knowledge may sufficiently advance to support prophylactic gene therapy to prevent polygenic late-onset diseases. A combination of a large number of common low effect gene variants offers the most likely explanation for the heritability of the majority of these diseases, and their risk may be lowered by correcting the effect of a subset of such gene variants.

**Methods:** Computer simulations quantified the correlation between the aging process, polygenic risk score, and hazard ratio change with age, using as inputs clinical incidence rates and familial heritability, and estimated the outcomes of hypothetical future prophylactic gene therapy on the lifetime risk and age of onset for eight highly prevalent late-onset diseases.

**Results:** The simulation results confirmed that gene therapy would be beneficial in both delaying the age of onset and lowering the lifetime risk of the analyzed lateonset diseases. Longer life expectancy is associated with a higher lifetime risk of these diseases, and this tendency, while delayed, will continue after the therapy.

**Conclusions:** When the gene therapy as hypothesized here becomes possible, and if the incidences of the treated diseases follow the proportional hazards model with multiplicative genetic architecture composed of a sufficient number of common small effect gene variants, then (**a**) late-onset diseases with the highest familial heritability will have the largest number of variants available for editing, (**b**) diseases that currently have the highest lifetime risk, and particularly those with the highest incidence rate continuing into older ages, will prove the most challenging cases in which to lower lifetime risk and delay the age of onset at the populational level, and (**c**) diseases that are characterized by the lowest lifetime risk will show the strongest and longest-lasting response to such therapy.

## Introduction

In the past two decades, the human genome was successfully sequenced. Whole genome sequencing (WGS) and genome-wide association studies (GWASs) of human and other organisms’ genomes have become an everyday occurrence (Visscher et al., 2017). Our knowledge of genetic variants, particularly the single nucleotide polymorphisms (SNPs) associated with susceptibility to diseases, has become deeper and more extensive.

The experimental gene therapy techniques aimed at diseases caused by a single defective gene or a single SNP— the so-called Mendelian conditions—are being refined. Mendelian conditions cause high mortality and morbidity, but each of these conditions affects only a minute fraction of the population. As of November 2018, the OMIM Gene Map Statistics compendium listed almost 6,300 phenotypic genetic conditions caused by close to 4,000 gene mutations. This list includes a variety of conditions, with onsets ranging from very early to late. For example, type 1 diabetes mellitus is caused by single defects in the HLADQA1, HLA-DQB1, or HLA-DRB1 genes (Steck and Rewers, 2011). Early-onset Alzheimer’s Disease (EOAD) is caused primarily by APP, PSEN1, or PSEN2 gene mutations and affects a relatively small proportion of the population, starting in their thirties, with the majority of mutation carriers being affected by the age of 65 (Ghani et al., 2018). In contrast, macular degeneration (Jager et al., 2008; Manolio et al., 2009; Sobrin et al., 2012) is primarily caused by a small number of high-effect variants and manifests at a relatively old age. In some cases, individualized genetic diagnoses—where an SNP that needs to be edited can be specified precisely—are possible. Over the last two decades, 287 monogenic disease clinical trials have been conducted worldwide (Ginn et al., 2018). When the medical technology becomes available, individuals who receive treatment will be effectively cured and will have no need for concern about the single, specific cause of their disease.

Polygenic or complex late-onset diseases (LODs) pose a more nuanced problem, and this study will focus on them. There are thousands of estimated gene variants or SNPs of typically small effect that, in combination, constitute the polygenic LOD risk of each individual (Eyre-Walker, 2010; Yang et al., 2012). These diseases include the old-age diseases that eventually affect most individuals and are exemplified by cardiovascular disease—particularly coronary artery disease (CAD)—cerebral stroke, type 2 diabetes (T2D), senile dementia, Alzheimer’s disease (AD), cancer, and osteoarthritis.

What distinguishes polygenic LODs from infectious diseases or from Mendelian genetic conditions is difficulty with the concept of *cure*. The diseases of aging are primarily a consequence of an organism’s decline over time, leading to increased susceptibility to many LODs (Ribezzo et al., 2016; Nelson and Masel, 2017; Fedarko, 2018). The combination of genetic liability, environmental factors, and the physiological decline of multiple organ systems leads to individual disease presentations (Franceschi et al., 2018). Detrimental gene variants are exacerbating factors (Gonzaga-Jauregui et al., 2012), compared to the average distribution of common gene variants that define human conditions as they apply to polygenic LODs. The time of onset for each individual is modulated by genotype and environment (Yashin et al., 2016). While some individuals will be diagnosed at a relatively young age, others will not be diagnosed with a particular LOD during their lifetime (Rzhetsky et al., 2007). According to the current consensus, a large number of common low-effect variants offer the likeliest explanation for the heritability of the majority of complex traits (Anderson et al., 2011; Yang et al., 2015). For example, in the cancers analyzed in this study, the fraction of all diagnoses that are attributed to highly detrimental inherited mutations is relatively low; it is estimated to explain 10–14% of breast cancer heritability (Risch et al., 2006; Haley, 2016), 10–12% of prostate cancer heritability (Wu and Gu, 2016; Mancuso et al., 2016; Walsh, 2017; Lecarpentier et al., 2017), and 5–10% of colorectal cancer heritability (de Voer et al., 2016; Graff et al., 2017) and is assumed to be a relatively minor fraction for lung cancer (Weissfeld et al., 2015; Wang and Wang, 2017; Kanwal et al., 2017). For the majority of these cancers, liability is attributed to the common low-effect gene variants and environmental factors. The development of cancer is a multistage process wherein individual variability in any tumorigenesis stage duration or liability may be influenced by hereditary predisposition as well as environmental factors (Weinberg, 2013). The level of susceptibility to the major polygenic LODs, and the difference between high-risk and low-risk individuals, may lie in a slightly higheror lower-thanaverage fraction of detrimental gene variants.

Certainly, the failure does not begin immediately prior to the age of diagnosis. For example, AD deterioration begins decades before symptoms first become noticeable (Alzheimer’s Association, 2017). A similar situation holds for cardiovascular disease (Lakatta and Levy, 2003; Lakatta, 2015) and cancer (Mitchell et al., 2018).

The best cure is prevention, and the time may be nearing when prophylactic gene therapy will be attempted for the prevention of complex polygenic diseases. Many questions of scientific knowledge, technical expertise, and ethics will need to be settled before this level of prophylactic gene therapy can become possible. From the ethical perspective, as techniques develop and the medical possibilities offered by gene therapy for improving health and preventing diseases gradually materialize, its acceptance is becoming more widespread. This is exemplified by the findings of the US Committee of the National Academies of Sciences, Engineering, and Medicine (2017) titled *Human genome editing—Science, ethics, and governance*, and the recommendations of the UK Nuffield Council on Bioethics (2018) *Genome editing and human reproduction: Social and ethical issues*, which considers germline editing as one possible application.

Computational techniques attempting to evaluate the effects of mutations or gene variants are being developed, although their accuracy needs to improve dramatically before they can become applicable to personalized human genetic evaluation or treatment (Liu et al., 2016). Similarly, while there are extensive libraries of human SNPs, including dbSNP, HapMap, SNPedia and aggregating sites (Beck et al., 2014), the information is far from actionable as far as modifying multiple personalized SNPs is concerned. The ability to locate or be able to computationally estimate a complete set of the low-effect causal SNPs requires knowledge that may take decades to gain. Gene editing technologies may also be a few decades away from the time when they can be used as routinely and with the same low risk as applying an influenza vaccination to modify a large number of gene variants distributed across the human genome. The latest gene editing technique, CRISPR-Cas9 (Wright et al., 2016), has supplemented and mostly replaced older technologies such as zinc-finger nuclease (ZFN) (Carroll, 2011) and transcription activator-like effector nuclease (TALEN) (Joung and Sander, 2013), though, for some applications, these older techniques continue to be more appropriate. While its selectivity and on-target precision are improving, CRISPR is still most effective in gene knockdown operations. For modification and repair, only a small fraction of CRISPR operations, using homologous repair with a template or a sister chromatid sequence, succeed. Nevertheless, CRISPR, which is only five years old, remains a rapidly developing technology that holds great promise. Synthetic genomics (Thompson et al., 2017; Kohman et al., 2018) could be another promising future technology. Synthetic genomics techniques could also help in developing the precise mapping of the effects of gene variants on disease phenotypes. If none of these approaches ultimately succeed in becoming reliable enough for the purposes of gene therapy, it is almost certain that a new, more suitable technique will be invented.

Changes in lifestyle and medical care, including the prevention and treatment of infectious diseases, have extended longevity over the last century, and this trend is projected to continue. This increased longevity is partly due to medical advances helping people live and function decades after first being diagnosed with historically deadly or debilitating illnesses. Preventive gene therapies may also become a future factor in prolonging healthspan. Actuarial science has tracked human mortality trends for centuries. The Gompertz–Makeham law of mortality, which was established more than 150 years ago, depicts an exponential increase in the rate of human mortality after the age of 30 (Makeham, 1860; Missov and Lenart, 2013). While the parameters of the Gompertz–Makeham law continue to be adjusted, the principle remains valid. The apparent squaring of the mortality curve—the so-called compression of morbidity and mortality into older ages—implies that the maximum human lifespan is likely limited to about 120 years of age (Stallard, 2016; Geddes, 2016; Barbi et al., 2018).

Within the next few decades, gene therapy techniques and genetic knowledge may sufficiently advance to support prophylactic gene therapy to prevent late-onset diseases. It may be timely to evaluate the extent of the effects that future gene therapies may have on delaying the onset of LODs or preventing them entirely.

The goal of this study was to establish how the proportional hazards model and multiplicative genetic architecture can be used to map the polygenic risk to hazard ratio of succumbing to the common late-onset diseases with advancing age and apply this mapping to quantify the effects of a hypothetical future prophylactic gene therapy. This study used as its foundation earlier research (Oliynyk, 2018), which reviewed epidemiology, heritability, and polygenic risk models, and developed a simulational basis for the analysis of eight of the most common diseases: AD, T2D, cerebral stroke, CAD, and breast, prostate, colorectal, and lung cancers. Computer simulations in this study quantified the correlation between the aging process, the polygenic risk score (PRS), and the change in the hazard ratio with age, using as inputs the clinical incidence rate and familial heritability, and estimated the outcomes of hypothetical future prophylactic gene therapy on the lifetime risk and age of onset for these eight LODs, and to estimate the lifetime risk increase associated with longevity gains.

## Methods

### Conceptual summary of the earlier study

This section briefly summarizes the concepts of the earlier research (Oliynyk, 2018) used as a foundation for this study. The subsequent sections will describe this study’s simulation flow and implementation.

### Cox proportional hazards model

According to Chatterjee et al. (2016), the conditional agespecific incidence rate of the disease, *I* (*t |G*), defined as the probability of developing the disease at a particular age *t*, given that a subject has been disease-free until that age, can be modeled using Cox’s proportional hazards model (Cox, 1972):

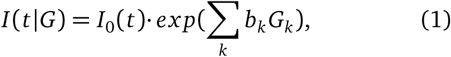

where *G* = (*G*_1_,…, *G*_*k*_) is the multiplicative effect of a set of risk factors on the baseline hazard of the disease *I*_0_(*t*). The set of age-independent variables in G could include genetic and environmental risk factors, as well as their interaction terms.

According to Chatterjee et al. (2016), if it can be assumed that environmental risk factors are independent of the SNPs, the *“post-GWAS epidemiological studies of gene-environment interactions have generally reported multiplicative joint associations between low-penetrant SNPs and environmental risk factors, with only a few exceptions.”* This means that the polygenic score *G* =Σ_*k*_ *b*_*k*_*G*_*k*_, as the lifelong characteristic of each individual, is used multiplicatively with *I*_0_(*t*), which encompasses environmental and aging effects.

This research’s simulations operate on model genetic architectures of the analyzed LODs, not a complete GWAS map of their experimentally discovered SNPs, because GWAS-discovered sets can explain only a fraction of these LODs’ heritability. These model genetic architecture SNPs are treated as “true” causal for disease liability and heritability variants, as discussed in Chatterjee et al. (2016), rather than GWAS-linked SNPs, and it is assumed that they can be accurately identified for the purposes of personalized gene therapy.

### Allele distribution models

The allele distribution models were based on (Noh et al., 2006; Pawitan et al., 2009) and are summarized next. An individual polygenic risk score *β* can be calculated as a sum of all alleles’ effect sizes, which is by definition a log(OR) (natural logarithm of odds ratio) for each allele:

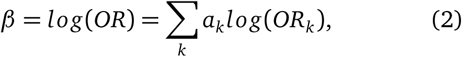

where *a*_*k*_ is the number of risk alleles (0, 1 or 2) and *OR*_*k*_ is the odds ratio of additional liability presented by the kth allele. Variance in the allele distribution is determined by:

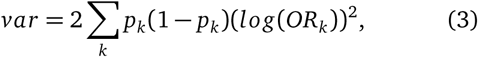

where *p*_*k*_ is the frequency of the k-th genotype (Pawitan et al., 2009). The contribution of genetic variance to the risk of contracting the disease is heritability:

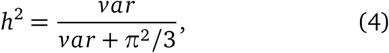

The *common allele low effect size* genetic architecture model (Scenario A, Table 1) is expected to be most suitable for explaining the heritability of the analyzed LODs, and all results are reported using this scenario. For verification, all simulations were also performed using a more extreme *rare allele medium effect size* model (Scenario B, Table 1), and the results for lifetime risk were essentially identical to those in Scenario A. Any material difference in the results would have warranted additional investigation. The number of variants needed for the Scenario A LODs is summarized in Table 2.

**Table 1.** Genetic architecture scenarios with modeled allele frequencies and effect sizes

| Scenario | MAF range | OR range | MAF values | Allele OR values |
| --- | --- | --- | --- | --- |
| A. Common low | 0.073-0.499 | 1.05-1.15 | 0.073 0.18 0.286 0.393 0.5 | 1.05 1.075 1.1 1.125 1.15 |
| B. Rare medium | 0.0146-0.0998 | 1.28-2.01 | 0.0146 0.036 0.0572 0.0785 0.0998 | 1.28 1.463 1.645 1.828 2.01 |

To build the genetic architecture, minor allele frequencies (MAFs) and ORs are chosen using 25 possible combinations of the values in the table. Following Pawitan et al. (2009), the variants are assigned to individuals with frequencies proportionate to MAF $p_k$ for SNP $k$ , producing, in accordance with the Hardy–Weinberg principle, three genotypes (AA, AB or BB) for each SNP with frequencies $p_k^2$ , $2p_k(1 - p_k)$ and $(1 - p_k)^2$ , resulting in the normal distribution of individual risk ORs.

**Table 2.**
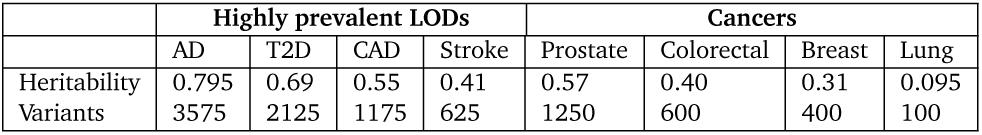
Heritability of analyzed LODs and an example of required variant numbers for common low-effect variants (scenario A)

|  | Highly prevalent LODs |  |  |  | Cancers |  |  |  |
| --- | --- | --- | --- | --- | --- | --- | --- | --- |
|  | AD | T2D | CAD | Stroke | Prostate | Colorectal | Breast | Lung |
| Heritability | 0.795 | 0.69 | 0.55 | 0.41 | 0.57 | 0.40 | 0.31 | 0.095 |
| Variants | 3575 | 2125 | 1175 | 625 | 1250 | 600 | 400 | 100 |

The allele scenarios are reused here for convenience as they were already implemented and validated in the earlier study and originally validated by Noh et al. (2006) and Pawitan et al. (2009). This study’s simulation does not need to track individual alleles, and the use of a normal effect size distribution with variance calculated using rearranged Eq. (4) and normalized to the population mean would produce the same distribution.

### LOD incidence rate functional approximation

The incidence rate functional approximations and source data are detailed in Oliynyk (2018). The yearly incidence rate logistic and exponential regressions for the functional approximation of the LODs based on the available clinical incidence statistics are illustrated in Fig. S4 in Supplementary File 1. The logistic approximations are used for all LODs except breast cancer, for which the exponential followed by linear regression more accurately approximates the incidence rate pattern.

### The aging coefficient: mapping PRS to age dependent probability of LODs

The earlier research (Oliynyk, 2018) showed that the incidence rates for these LODs increase as individuals age, and statistically, the PRSs of individuals diagnosed at older ages decline. Following the multiplicative model of PRS and environmental effects, from a statistical perspective, two processes are balancing disease incidence: The aging or environmental effect increases the average susceptibility of the population, and this makes it more likely that older individuals will become ill. Individuals with higher PRSs are more likely to become ill, and with higher incidence rates, a larger fraction of these individuals have already become ill with every additional year of age and are therefore excluded from the high-risk pool for the following years of age. The aging process continues, as reflected in the incidence rate rising or remaining high in subsequent years.

Statistically, individual PRSs and environmental effects, including aging, affect the probability of a person becoming ill (Chatterjee et al., 2016). According to Cox’s proportional hazards model (Cox, 1972), in every year of age, each individual can be assigned a hazard ratio which describes that individual’s risk of being diagnosed with an LOD. The goal is to uncover the mapping between PRSs and individual hazard ratios on a yearly basis:

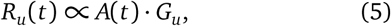

where *R*_*u*_(*age*) is the hazard ratio of *u*th unaffected individual, and *t* is the individual’s age in years. PRS *G*_*u*_ remains constant for each individual for life, and the multiplier *A*(*t*) drives the age-related increase in LOD incidence. *A*(*t*) will also be called the “aging coefficient”.

It may be possible to map or “discover” *A*(*t*) by applying the yearly incidence rate in a population to the PRS distribution, based on an LOD genetic allele architecture, through the population simulation flow described in the next section. The discovered *A*(*t*), then applied to the population simulation on a yearly basis, should precisely reproduce the LOD initial incidence rate pattern. Later, building a population with a modified PRS to emulate the effect of gene therapy, and simulating aging of this population by applying *A*(*t*), it will be possible to find the corresponding incidence rate and other resulting statistics. In addition to the age-related change in PRS distribution described above, the aging coefficient automatically incorporates other miscellaneous environmental effects that accumulate with age, because these are all reflected in the yearly incidence rate. The term “aging coefficient” is used throughout this publication rather than “environmental effect” to emphasize that it is an aggregate age-dependent parameter.

A useful parameter in the simulation and analysis is incidence rate density *D*(*t*), which depicts LOD incidence contributions relative to the initial population count at yearly increments, adjusted for mortality:

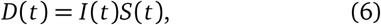

where *I* (*t*) is the yearly incidence at age *t* and *S*(*t*) is the survivor rate from the US Social Security Actuarial Life Table. Integrating the area under the curve, or summing up discrete yearly values, corresponds to an LOD cumulative incidence *C* (*t*); the limit or lifetime value *C* (*T)* equals lifetime risk:

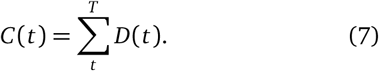

### Simulation steps in discovering LOD aging coefficients

1. The simulation initialization steps are performed, including allocating population objects, assigning individual PRSs based on the modeled genetic architecture, and building an incidence rate functional approximation.
2. The simulation works as an iterative procedure, where the values of *A*(*t*) are matched at advancing ages, starting at the age at which the incidence rate first becomes noticeable for an LOD, denoted as *t* = *T*_0_. The initial value is relatively immaterial, and each simulation begins with the value *A*(*T*_0_) = 0.001, The iterative process, by following steps 3 and 4, rapidly finds a close match value. For instance, in the case of AD, age *T*_0_ = 39 years, *I* (39) = 2.06·10^*-*6^, and the resulting *A*(39) = 5.85·10^*-*9^. *A*(*t*) discovery for the next year commences with the value for the previous year and as a result requires slightly fewer iterations to find the match.
3. As the *A*(*t*) value is applied to all individuals’ PRSs, a first estimate of each individual hazard ratio for this year is produced. Based on the resulting probabilities, a number of individuals will be diagnosed at age *t*. This number is simply *N*_*d*_ (*t*) = *I* (*t*) *N*_*u*_(*t*), where *N*_*u*_(*t*) is the remaining healthy population and *N*_*d*_ (*t*) is the number of individuals expected to be diagnosed.
4. The simulation verifies how well the result matches the expected incidence rate for the LOD at this age, and a better-matching approximation value is recalculated and reapplied to step 3. This process will iterate steps 3 and 4 until a predetermined level of accuracy is achieved, with the aim of attaining 0.1% reproduction accuracy for ages with significant incidence rates.
5. This cycle of steps 3 and 4 is then rerun a predetermined number of times (with 10 repetitions being the default), with iterations alternatively commencing with a value *A*(*t*) 10% above and below the previously determined value to account for a potential determination bias. The results of these reruns are averaged, and the resulting variance is evaluated and recorded.
6. The validated aging coefficient *A*(*t*) from step 5 is then applied to the step 3 operation one final time in order to sample the individuals diagnosed at this age out of the population. Statistically, the highest-PRS individuals are likelier to be diagnosed, and these individuals are excluded from the healthy population.
7. The age is advanced by one year, with a now-smaller population *N*_*u*_(*t* + 1) = *N*_*u*_(*t*) − *N*_*d*_ (*t*). The simulation repeats steps 3–6 until all ages are covered—in our case, until the age of 120.

At completion, the aging coefficients that map the modeled PRS value to the individual hazard ratio for each year of age have been discovered.

### Calculating cumulative incidence and lifetime risk for discrete LOD hazard ratios

The cumulative incidence is determined by applying the aging coefficients found in the discovery simulation above to a range of hazard ratio (HR) multiples relative to the population mean. As reported in the Results section, these calculations allow for the evaluation of how cumulative incidence and lifetime risk correspond to hazard ratios; this is equivalent to learning what would happen if the individual hazard ratio were changed using gene therapy (or, in principle, any intervention of a similar effect). An R script (available in Supplementary Data 1) calculates the incidence for a fixed value of hazard ratio *G*_*k*_ through the range of ages from 0 to 120:

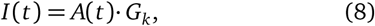

and Eq. (7) gives the age progression of lifetime risk. The HR values used were: 16.0, 4.0, 1.0, 0.25 and 0.0625; the script could be modified for any set of desired values.

Here, an R script takes the aging coefficient as the input and calculates the lifetime risks for PRS = 1.0 (population mean risk), PRS = 0.5, and PRS = 0.25 and the numbers of years by which life expectancy must increase in order for the lower PRS value to again exceed the mean population lifetime risk, reported in the Results section for fourfold PRS changes: 16.0 *→* 4.0, 4.0 *→* 1.0, and 1.0 *→* 0.25.

### Simulating outcomes of gene therapy lowering population polygenic risks

In this simulation, the aging coefficient discovered through the earlier simulation is applied while the population ages, and the resulting population incidence rate and lifetime risk patterns are analyzed.

It is possible to conceive any number of scenarios of prophylactic gene therapy. In the most extreme scenario, one could make an equivalent of thousands of edits for AD, and hundreds for most of the rest of the analyzed LODs, reversing all variants constituting individual PRSs to a neutral state; this would show a resulting disease risk and incidence of zero. It is likely that any models featuring so drastic an intervention would be unrealistic, and many less extreme scenarios are possible.

The simulation scenario where every population’s individual PRS is decreased fourfold (OR=0.25) was chosen as a sufficient number of edits to achieve a substantial effect: It delays the lifetime risk of T2D, stroke and CAD by almost precisely 15 years and provides even better outcomes for all analyzed cancers, while highlighting the difficulty of treating AD. A life extension of 15 years would result in an average life expectancy of about 95 years, and this may be considered as challenging the unknown limits of the squaring of the mortality curve sufficiently far for this study.

It appears most practical for a person to be born with all cells already treated, because all developmental stages are affected by the genome, which implies germline therapy and likely heritability of therapy. Nevertheless, it is not yet known what technological possibilities and ethical considerations the future will bring; therefore, without discussing the specifics of any method, it is assumed that, at birth (or age zero), the person’s genome appears with the required modification. The therapy emulation is a simple arithmetic operation of reducing the individual PRS by a desired value. Simulating a fourfold reduction in all population individuals’ PRS (OR = 0.25, equivalent to *β* = log(OR) = −1.39) requires an edit of 15 SNPs in a common allele low-effect-size scenario with an average OR of 1.1; it is simply calculated as: 1.1^15^*≈* 4. Choosing to edit only the largest-effect-size SNPs available in the genetic architecture, with OR = 1.15, would require approximately 10 edits: 1.15^10^≈4. The simulations were then performed with the previously discovered values of *A*(*t*) and the population with the modified individual PRSs.

The lowerand higherintensity scenarios were also simulated and were qualitatively similar to the simulation above, with a corresponding decrease or increase in lifetime risk and onset delay patterns; the above scenario was found to be the most representative and illustrative and is reported in the Results section. A somewhat different scenario, in which all individuals with elevated PRSs were treated to adjust their PRSs to that of the population mean, merited an illustration in Fig. S5 in Supplementary File 1 and Fig. S6 in Supplementary File 1; the lifetime risk outcomes are also qualitatively similar to those in the above simulation.

### Emulating life expectancy increases

Life expectancy increases are emulated by adjusting the mortality from all causes by the desired number of years. All mortality rates from the US Social Security Actuarial Life Table are shifted by 5, 10, and 15 years. This approach is supported by Zuo et al. (2018), who showed that the front slope between the 25th and 90th percentiles of old-age deaths advanced with a nearly constant long-term shape as longevity increased over the past five decades. It may be not prudent to heed the opinion expressed in Zuo et al. (2018) that there is “no support for an impending limit to human lifespan” indefinitely, but it may be sufficient for this study’s estimates of up to 15 years of increased life expectancy. Although the Results section reports values with greater life expectancy increases, these have primarily comparative value, and *>* 40 years will denote even greater extensions.

### The aging coefficient values and simulation accuracy

Fig. 1 shows the resulting accuracy produced by applying the aging coefficient to eight LODs, compared to the input incidence rate approximation. The error rate begins near 2% for colorectal and lung cancers, is noticeably better than 1% for the remaining LODs, and stays below 0.2% for most of the LODs’ onset range. This validation illustrates that the combined discovery and analysis reproduce the input LOD incidence rate with a high degree of precision.

**Figure 1.**
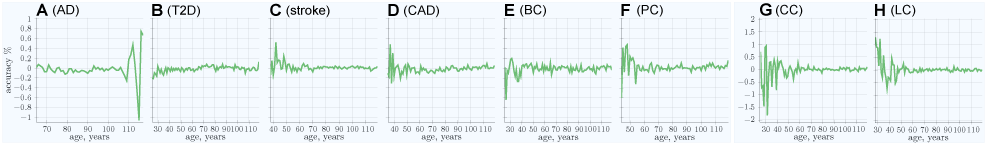
Accuracy of simulation aging coefficient discovery in reproducing the input incidence rate approximation. **(A)** Alzheimer’s disease, **(B)** type 2 diabetes, **(C)** cerebral stroke, **(D)** coronary artery disease, **(E)** breast cancer, **(F)** prostate cancer, **(G)** colorectal cancer, **(H)** lung cancer.

The core of the simulation is the iterative discovery and application of the aging coefficient (see Methods Eq. (5)), mapping individual PRSs to the hazard ratio specific to each LOD on a yearly basis, depicted in Fig. 2. The aging coefficient incorporates combined aging and environmental effects, and the rising pattern indicates the increasing magnitude of these effects with age. Fig. 3 shows the PRS distribution for diagnosed individuals by age when the simulation is rerun using the discovered aging coefficient without gene editing (a baseline validation). This distribution matches the results of Oliynyk (2018), in which the simulation used a direct probabilistic algorithm. A more precise indication of the accuracy of the discovered aging coefficient is shown in Fig. 1, where the simulation aging reproduces the population incidence rate Eq. (5).

**Figure 2.**
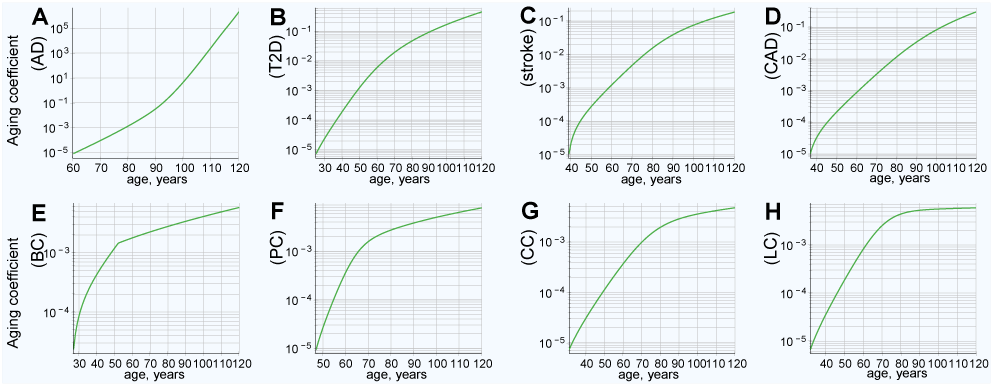
Aging coefficients reflects increases in LOD liability with age based on the clinical incidence rate and model genetic architecture PRS. **(A)** Alzheimer’s disease, **(B)** type 2 diabetes, **(C)** cerebral stroke, **(D)** coronary artery disease, **(E)** breast cancer, **(F)** prostate cancer, **(G)** colorectal cancer, **(H)** lung cancer.

**Figure 3.**
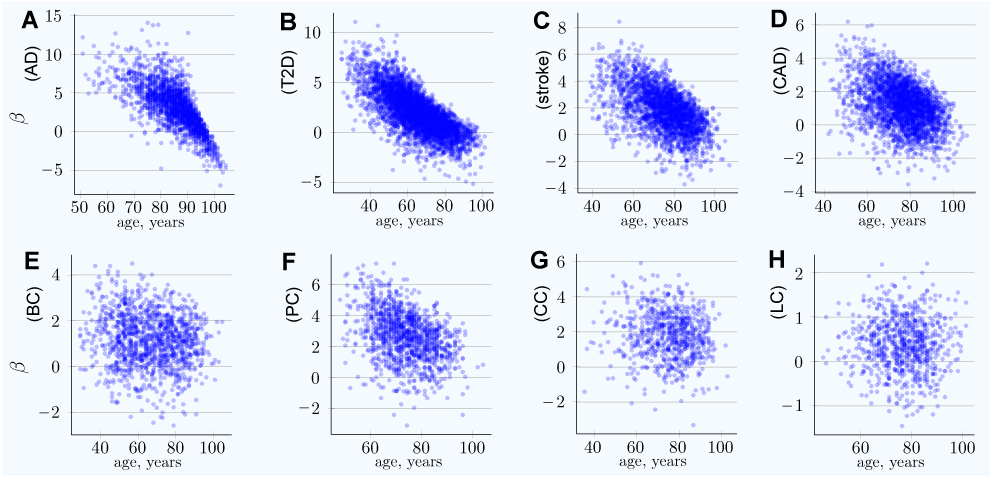
Aging coefficients applied to reproduce the LOD incidence distribution by age based on individual PRS. **(A)** Alzheimer’s disease, **(B)** type 2 diabetes, **(C)** cerebral stroke, **(D)** coronary artery disease, **(E)** breast cancer, **(F)** prostate cancer, **(G)** colorectal cancer, **(H)** lung cancer. Polygenic scores of individuals diagnosed with an LOD as a function of age. Scatter plots show the distributions of polygenic scores for cases diagnosed as age progresses. *PRS β* = *log*(*OddsRatio*).

The values with lower accuracy occur at very early and very late ages of onset, affecting a minute fraction of the total population. This in itself is the cause of the variability. As seen in Fig. S1 in Supplementary File 1, the baseline case, and in Fig. S2 in Supplementary File 1, the gene therapy case, the incidence rate density is very low at younger ages, when it is below 1· 10^*-*5^. Due to population mortality, even though the incidence rate is high at old ages, the number of individuals still alive after the age of 105 is small, even with an increase in the modeled life expectancy of 15 years. Only in the case of AD, the incidence of which grows exponentially to a very late age, does the remaining unaffected population become small near 120 years of age, causing the error rate to increase to close to 1%, a minuscule deviation in light of the particularly low incidence rate density at this age.

This precision is due to the use of large population sets: 25 million simulated individuals, with aggregation of 250 simulation loops (configurable), for each simulation iteration in the discovery stage and a population of one billion for the gene therapy simulation. While the analysis could certainly have been performed with smaller simulation sets, leveraging the available computing equipment allowed for the achievement of low statistical variance while simplifying analysis. This precision rendered the use of error bars in the graphical displays impractical.

### Data sources, programming, and equipment

The population mortality estimates from the US Social Security Actuarial Life Table provide annual death probability and survivor numbers up to 119 years of age for both men and women.

Disease incidence data were obtained from the following sources: Alzheimer’s disease (Brookmeyer et al., 1998; Edland et al., 2002; Kokmen et al., 1988; Hebert et al., 1995), type 2 diabetes (Boehme et al., 2015), coronary artery disease and cerebral stroke (Rothwell et al., 2005), and cancers (Cancer Statistics for the UK; Kuchenbaecker et al., 2017).

The simulations were performed on an Intel Xeon Gold 6154 CPU-based 36-core computer system with 288GB of RAM. The simulation is written in C++, and the source code can be found in Supplementary Data 1.

The final simulation data, additional plots, R scripts, and executables are also available in Supplementary Data 1. Intel Parallel Studio XE was used for multi-threading support and Boost C++ library for faster statistical functions; the executable can be built and function without these two libraries, with a corresponding slowdown in execution.

## Results

### Characteristics of the aging coefficient

The clinical incidence rate pattern was used to map the hazard ratio from the PRS of individuals diagnosed with an LOD at each year of age for eight LODs: AD, T2D, cerebral stroke, CAD, and four late-onset cancers: breast, prostate, colorectal, and lung. Knowledge of the hazard ratio for any given PRS value at any given age—the aging coefficient—is what is needed if the further calculations and simulations are to model gene therapy and the effects of life expectancy increases. The aging coefficient was discovered for these LODs, as described in the Methods section “Aging coefficient discovery simulation steps,” and can be seen in Fig. 2.

It is interesting to note how the magnitude of the aging coefficient changes with age for the analyzed diseases. The range of values spanned by the aging coefficient and its maximum value, seen in Fig. 2A, is larger for AD than for all other analyzed LODs. This can be ascribed to two factors that distinguish AD: (1) the steepest rise of incidence rate and cumulative incidence, and (2) the highest heritability of the reviewed LODs, resulting in the highest variance according to our genetic architecture model. This larger variance results in more extreme values of high and low PRSs in the population, and therefore the age coefficient multiplier is necessarily smallest at the youngest onset ages, with the opposite being true at the older ages. T2D, CAD, and stroke show a comparatively more moderate progression and maximum projected aging coefficient values, while cancers show a still-smaller maximum magnitude. Lung cancer possesses both the lowest heritability and the lowest cumulative incidence, and consequently the values discovered by simulation for the lung cancer aging coefficient (seen in Fig. 2H) almost precisely match its incidence rate. In a limit case of an LOD characterized by a PRS that remains constant with age and a very low incidence rate, the aging coefficient should be identical to the incidence rate.

### Longer life expectancy corresponds to increasing lifetime risk

The modeled increased life expectancy in the baseline scenario, without prophylactic gene therapy, is displayed in Fig. 4 and summarized in Table 6. This analysis corresponds to the baseline incidence rate represented by the blue line in Fig. 5. All analyzed LODs show a significant lifetime risk increase with every five years of life expectancy extension. This is most prominent with AD, the incidence of which nearly triples with an extension of life expectancy of 15 years, while the incidence rate less than doubles for the rest of the LODs. The incidence rate density (as seen in Fig. S1 in Supplementary File 1) shows the relative incidence increasing while the peak of incidence shifts toward older ages.

**Figure 4.**
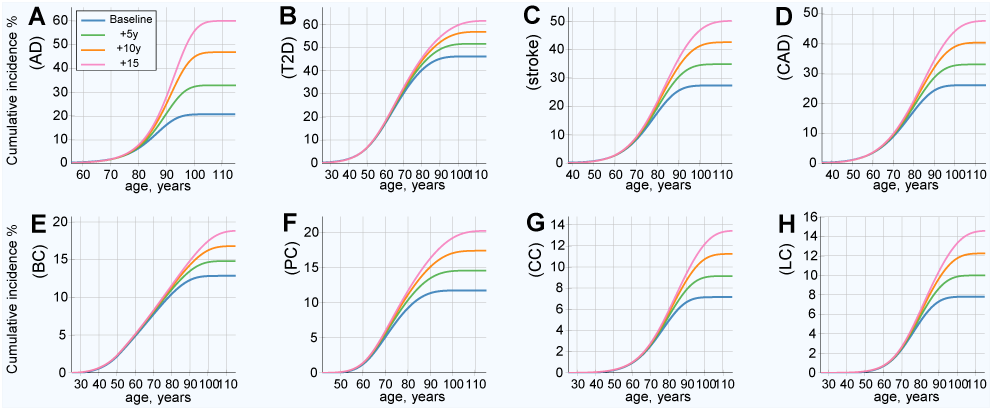
Projected LOD cumulative incidence and lifetime risk increase if life expectancy increases by 5, 10 and 15 years (baseline without gene therapy). **(A)** Alzheimer’s disease, **(B)** type 2 diabetes, **(C)** cerebral stroke, **(D)** coronary artery disease, **(E)** breast cancer, **(F)** prostate cancer, **(G)** colorectal cancer, **(H)** lung cancer. This is the baseline scenario without gene therapy or other health improvements for the plotted LOD. It represents the case where life expectancy increases due to causes other than the plotted LOD. Lifetime risk (lifetime cumulative incidence) corresponds to the lifetime (right-most) values of the plots.

**Figure 5.**
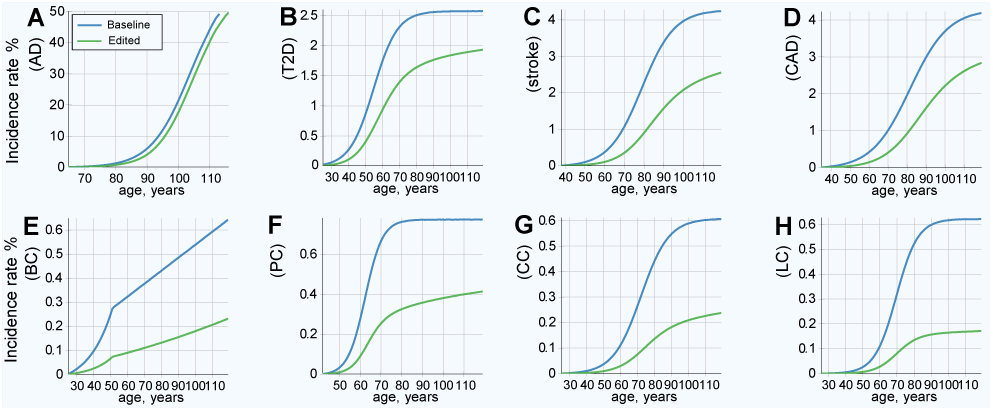
LOD incidence rate patterns, baseline and after emulated gene therapy. **(A)** Alzheimer’s disease, **(B)** type 2 diabetes, **(C)** cerebral stroke, **(D)** coronary artery disease, **(E)** breast cancer, **(F)** prostate cancer, **(G)** colorectal cancer, **(H)** lung cancer. All individuals in the population had emulated corrective gene therapy editing of, on average, 15 SNPs (corresponding to an OR multiplier of 0.25).

### Lifetime risk estimates for discrete hazard ratio multiples

Having the aging coefficient, it is easy to recalculate life-time incidence risk for a range of HRs. Population mortality is one of the principal limiting factors on the lifetime risk of an LOD, and shifting the mortality curve and thus emulating longer life expectancy reveals how incidence rate and lifetime risk would change if life expectancy were to increase.

The Fig. S3 in Supplementary File 1 shows a grid display with HR ranging from 16.0 to 0.0625 and average life expectancy varying from baseline to extended up to 15 years. The results of these calculations show that the lifetime risk is proportionate to the HR, as long as lifetime risk is relatively low (below 50%) for all reviewed LODs, even though they display quite varied incidence patterns. Lowering the HR through gene therapy or other means implies proportionately lowering the lifetime risk, assuming life expectancy remains constant, as summarized in Table 3. For instance, lowering the HR fourfold is accompanied by a fourfold, or 400%, drop in lifetime risk. The only exception is AD, for which the lifetime risk decreases at a slower-than-proportionate rate; a possible explanation could be the near-exponential incidence rate increase to very advanced ages for AD, while all other LODs an-alyzed here can be approximated by flattened or linear progression at a more moderate old age (Oliynyk, 2018). Fig. S3 in Supplementary File 1 shows a dramatic increase in the lifetime risk of LODs with every five extra years of life expectancy. Evidently, with increased life expectancy, the projection approaches certainty for higher HRs and increasing life expectancies. This is most prominent for AD and T2D and can be explained by the high heritability and high later-age prevalence of these LODs, leading to a significantly higher risk that individuals will become ill earlier. Furthermore, at an advanced age, the remaining lower-risk individuals are those who constitute the majority of incidence cases (Oliynyk, 2018). As a result, the modeled high-PRS individuals show improvement in delaying the statistical disease onset, while the lifetime risk may remain almost as high.

**Table 3.** Lifetime risk corresponding to select discrete LOD PRSs (%)

| PRS value | Highly prevalent LODs |  |  |  | Cancers |  |  |  |
| --- | --- | --- | --- | --- | --- | --- | --- | --- |
|  | AD | T2D | Stroke | CAD | Breast | Prostate | Colorectal | Lung |
| 1.0 | 43.3 | 78.7 | 26.8 | 21.1 | 8.27 | 3.94 | 3.33 | 7.12 |
| 0.5 | 31.9 | 39.9 | 13.5 | 10.6 | 4.14 | 1.97 | 1.66 | 3.56 |
| 0.25 | 22.0 | 20.0 | 6.75 | 5.30 | 2.07 | 0.984 | 0.832 | 1.78 |

The application of the aging coefficient and life expectancy increases makes it simple to estimate the onset delay if the PRS were changed, as in the case of prophylactic gene therapy. Tables 4 and 5 show the values of shifts in the onset delay on the cumulative incidence slope at 30% of lifetime risk and for the full lifetime risk, respectively. Emulating longer life expectancy raises lifetime risk, and we see a more complex picture than a mere proportionate decline in risk, depending on the incidence and heritability patterns for each LOD. The AD lifetime risk exceeded the baseline within *∼*1–4 years of longer life expectancy. It would take *∼*10–15 years of longer life for T2D, stroke, and CAD to approach or slightly exceed their respective baseline lifetime risks. In nearly all calculated scenarios, it took significantly longer than 15 years for cancers to exceed their baseline lifetime risks.

**Table 4.**
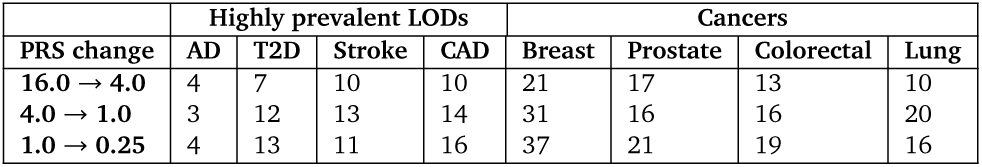
Estimate of LOD onset delay of incidence slope at 30% of lifetime risk (values in years)

| PRS change | Highly prevalent LODs |  |  |  | Cancers |  |  |  |
| --- | --- | --- | --- | --- | --- | --- | --- | --- |
|  | AD | T2D | Stroke | CAD | Breast | Prostate | Colorectal | Lung |
| 16.0 → 4.0 | 4 | 7 | 10 | 10 | 21 | 17 | 13 | 10 |
| 4.0 → 1.0 | 3 | 12 | 13 | 14 | 31 | 16 | 16 | 20 |
| 1.0 → 0.25 | 4 | 13 | 11 | 16 | 37 | 21 | 19 | 16 |

**Table 5.** Estimate of LOD onset delay for lifetime risk (values in years)

| PRS change | Highly prevalent LODs |  |  |  | Cancers |  |  |  |
| --- | --- | --- | --- | --- | --- | --- | --- | --- |
|  | AD | T2D | Stroke | CAD | Breast | Prostate | Colorectal | Lung |
| 16.0 → 4.0 | 1 | 0 | 2 | 3 | > 40 | 19 | 24 | 21 |
| 4.0 → 1.0 | 2 | 9 | 14 | 12 | > 40 | 28 | 34 | > 40 |
| 1.0 → 0.25 | 3 | 20 | 17 | 14 | > 40 | 29 | 35 | > 40 |

**Table 6.** LODs’ characteristics and lifetime risk in a range of life expectancy scenarios and with prophylactic gene therapy lowering population PRS fourfold

|  | Highly prevalent LODs |  |  |  | Cancers |  |  |  |
| --- | --- | --- | --- | --- | --- | --- | --- | --- |
|  | AD | T2D | Stroke | CAD | Breast | Prostate | Colorectal | Lung |
| Literature and clinical data: |  |  |  |  |  |  |  |  |
| Heritability | 0.795 | 0.69 | 0.55 | 0.41 | 0.57 | 0.40 | 0.31 | 0.10 |
| Literature max yearly incidence rate | > 20% | 2.5% | 4.4% | 3.6% | <0.5% | <0.8% | <0.6% | <0.6% |
| Lifetime risk, baseline + longer life: |  |  |  |  |  |  |  |  |
| +5 years life expectancy | 160% | 112% | 128% | 127% | 115% | 123% | 127% | 128% |
| +10 years life expectancy | 228% | 123% | 156% | 155% | 130% | 147% | 156% | 156% |
| +15 years life expectancy | 293% | 134% | 184% | 184% | 146% | 172% | 187% | 186% |
| Lifetime risk, OR 0.25 therapy vs baseline: |  |  |  |  |  |  |  |  |
| Therapy, unchanged life expectancy | 70% | 67% | 44% | 50% | 30% | 41% | 31% | 27% |
| Therapy, +5 years life expectancy | 124% | 77% | 61% | 69% | 36% | 52% | 40% | 34% |
| Therapy, +10 years life expectancy | 191% | 88% | 81% | 89% | 41% | 65% | 51% | 43% |
| Therapy, +15 years life expectancy | 260% | 98% | 101% | 112% | 47% | 77% | 62% | 52% |

LODs’ heritability and clinical incidence are discussed in the Methods section. The baseline is considered 100% in lifetime risk comparisons; 160% after a 5-year life expectancy extension for AD means an increase in lifetime risk (LR) of 1.6 times, and 50% after gene therapy means half the LR, compared to the baseline value.

Reviewing the first four LODs in the first row—and to a lesser extent AD and T2D in the second row—of Table 5, it is noticeable that, for high PRS values of 16.0 and 4.0, the response is lower than for the final row and also lower than for the slope onset delay in Tables 4. Fig. S3 in Supplementary File 1 shows that the likelihood of becoming ill is modeled as a near certainty for these values. The simulation results aggregating outcomes for the population in the following section provide additional generalization and confirmation of these patterns.

### Results of simulated gene therapy lowering population PRS

The results of this simulation show an improved (lower) incidence rate after treatment, as displayed in Fig. 5, and a corresponding decrease in lifetime risk, as presented in Fig. 6. Fig. 6 also serves as a good qualitative illustration of the results of the previous section. Comparing the baseline (blue line) to therapy with life expectancy (green line) held constant shows a significant improvement in both the lifetime risk and age of onset for all LODs. The lifetime risk is lower for all LODs and shows a delay of approximately a decade in the incidence rate curve for T2D, stroke, and CAD. AD benefits the least, and cancers show the most significant improvement. Table 6 illuminates the results detailed earlier from a slightly different perspective. With life expectancy unchanged, the lifetime risk decreases by 30% or more for AD and T2D, by more than 66% for colorectal and lung cancer, and 50% or more for the remaining four LODs.

**Figure 6.**
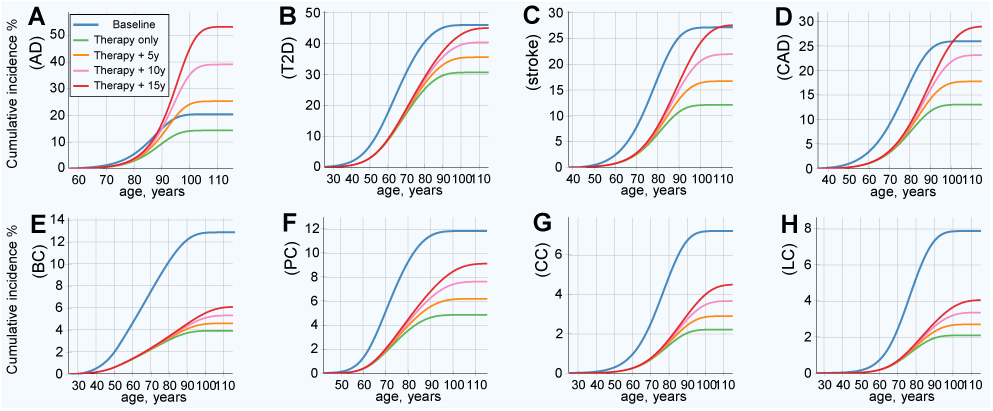
Cumulative incidence and lifetime risk following therapy reducing the population risk fourfold. **(A)** Alzheimer’s disease, **(B)** type 2 diabetes, **(C)** cerebral stroke, **(D)** coronary artery disease, **(E)** breast cancer, **(F)** prostate cancer, **(G)** colorectal cancer, **(H)** lung cancer. All individuals in the population had emulated corrective gene therapy editing of, on average, 15 SNPs (corresponding to an OR multiplier of 0.25). Result of gene therapy and results in increases in life expectancy of 5, 10, and 15 years. Lifetime risk (lifetime cumulative incidence) corresponds to the lifetime (rightmost) values of the plots.

Emulating longer life expectancy results in increased lifetime risk. The AD lifetime risk exceeded the baseline within *∼*3 years of longer life expectancy. It took 15 years of longer life for T2D, stroke, and CAD to approach or slightly exceed their respective baseline lifetime risks. Nevertheless, even with longer life expectancy, the onset remains delayed compared to the baseline. All cancers remained far below their baseline lifetime risks, even with a 15-year longer life expectancy. The risk remained lower than the baseline for all analyzed cancers after an increase of 15 years in life expectancy; any LOD with a relatively high heritability and low prevalence should similarly benefit. The results are summarized in Table 6 along with relevant LOD statistics.

## Discussion

For the purposes of this hypothetical treatise, it was assumed that it is possible to precisely identify individual gene variants and their detrimental or beneficial effects and then use gene editing to modify a large number of detrimental variants. This hypothetical gene therapy model was applied to estimate what would happen to LOD progression as the population ages. Conceptually, gene therapy here does not consider additions of artificially designed genomic sequences, but rather only corrections made to typically low-effect heterozygous in-population gene variants—that is, a correction of a detrimental variant to a naturally occurring neutral state. For the sake of simplicity, the model used SNP distributions, though the same would apply, albeit with a higher degree of complexity, to gene therapy using other gene variant types.

This study does not evaluate potential obstacles due to pleiotropy, defined for the purposes of gene therapy as the possible negative effect on other phenotypic features of any attempt to prevent an LOD by modifying a subset of SNPs (Stearns, 2010; Paaby and Rockman, 2013). The high-risk individual PRS is caused by numerous variants. In this model, these are normally distributed in the population. There is a relatively small difference in the absolute number of detrimental alleles between the population average and higher-risk individuals. Arguably, for the purpose of personalized prophylactic treatment, it will be possible to select a small fraction of variants from a large set of available choices seen in Table 2 that do not possess antagonistic pleiotropy, or perhaps even select SNPs that are agonistically pleiotropic with regard to some of the other LODs.

Applying the modeled aging coefficient to evaluate the impact of longer life expectancy on lifetime risk confirms the long-standing observation that aging itself is the predominant risk factor for many late-onset diseases and conditions. The lifetime risk increase with age is most prominent for AD. In those countries with longer life expectancy, the lifetime risk of AD is usually higher, as is demonstrated by Wu et al. (2017) using the example of Japan. These results confirm once more that, if mortality from all causes is lower, resulting in a longer life expectancy, AD is an LOD that exhibits the most rapid rise in advanced-age prevalence. The Framingham General Cardiovascular Risk Score includes age as one of the major risk factors for stroke and CAD (D’Agostino Sr et al., 2008). Boehme et al. (2015) show a similar pattern for T2D, and the results of this study are in agreement.

The calculations applying the discovered aging coefficient to discrete hazard ratio values show a delay in onset incidence for all analyzed LODs. The lifetime risk decreases in proportion with a decrease in hazard ratio as long as the absolute value of lifetime risk remains low. With the introduction of an emulated life expectancy increase, the lifetime risk increases.

The gene therapy simulation scenarios analyzing population statistics show decreases in LOD incidence and delays in LOD onset. These simulations also show the increase in lifetime risk with emulated longer life expectancy. Such estimates may be important for evaluating population health and well-being and the potential financial impact on healthcare systems. This study’s estimates, based on the proportional hazards model and multiplicative genetic architecture using the aging coefficient, allowed for an estimation of these effects accounting for a model genetic architecture of the LODs, rather than a more simplistic calculation based primarily on the statistical shape of the incidence rate progression. In a study aptly titled “Projections of Alzheimer’s disease in the United States and the public health impact of delaying disease onset,” Brookmeyer et al. (1998) estimate that an intervention that achieves a twofold AD hazard ratio decrease would shift the exponential rise curve of AD by five years, leading in the long term to a twofold decline in the cumulative incidence and prevalence of AD when accounting for mortality. The simulation reflecting age-related change in PRS distribution demonstrates that the positive effect on the lifetime risk of AD would be significantly lower than projected by the above study in the case of preventative gene therapy. While AD emerges as one of the most difficult diseases to prevent, LODs with low cumulative incidence, such as cancer, exhibit enduring improvement under this model.

Even though in this study each LOD is analyzed independently, in practice, prioritizing certain LODs for preventative therapy could have a significant effect on other conditions not specifically targeted for treatment. For example, T2D is one of the diseases that cause the most co-morbidities, accelerating onset of cardiovascular and other diseases, sometimes by decades (Boehme et al., 2015). For this reason, preventative treatment of T2D could mean improvements in health or delays in the presentation of a range of LODs, either independently of or in addition to treating their specific gene variants.

## Conclusions

Computer simulations in this study mapped polygenic risk to the hazard ratio of being diagnosed with eight common LODs based on their known heritability and incidence rates, under the proportional hazards model and multiplicative genetic architecture. The resulting mapping— the aging coefficient—enabled the researcher to quantify the population effects of the emulated prophylactic gene therapy alongside longevity increases. Computer modeling and simulations deal with simplifications and generalizations of biological processes, and aim to make predictions about modeled systems behavior when modifying parameters of a model; the conclusions of this study are made in such context.

The intensive gene therapy simulated here could dramatically delay the average onset of the analyzed LODs and reduce the lifetime risk of the population. The simulations highlighted that the magnitude of familial heritability and cumulative incidence patterns distinguish the outcomes for the analyzed LODs when subjected to the same PRS decrease. This outcome can be characterized by the delay in LOD onset—the estimate of the number of years it would take for each LOD to regain the pre-treatment baseline level.

It would be difficult to limit the prevalence of AD, which is delayed only by *∼*3 years with the modeled level of therapy. AD may require a higher number of gene edits, likely postponing the possibility of more effective treatment to a point even further in the future; yet any improvement would be welcome. It is possible that a pharmaceutical intervention targeting a causal metabolic pathway or immune or inflammatory response may be more effective for AD, although the past announcements generating false hope regarding breakthroughs in these approaches are too numerous to cite.

For T2D, stroke, and CAD, lifetime risk will regain pretreatment baselines within 10 to 15 years of longer life, which is equivalent to delaying the average onset age of these LODs by as many years. Based on heritability and incidence rate combinations, prophylactic gene therapy holds the potential to bring significant and longer-lasting benefits for cancer prevention, even with a similar or smaller number of edited gene variants than for the more prevalent diseases. Additionally, when such advanced gene therapy technologies become available, preventing monogenic, highly detrimental variants will be simple, and the combination of therapies can bring about even more substantial improvements in both individual and populational health outcomes.

These conclusions can be generalized as follows: When gene therapy as hypothesized here becomes possible, and if the incidence of the treated diseases follow the proportional hazards model with multiplicative genetic architecture composed of a sufficient number of common low effect gene variants, then (**a**) late-onset diseases with the highest familial heritability will have the highest number of variants available for editing, (**b**) diseases with the highest current lifetime risk, and particularly with the highest incidence rate continuing into advanced age, will be the most resistant to attempts to lower the lifetime risk and delay the age of onset at the populational level, and (**c**) diseases that are characterized by the lowest lifetime risk will show the strongest and longest-lasting response to such therapy.

## Acknowledgments

The author thanks Alexei J. Drummond at the University of Auckland for a number of helpful and challenging discussions.

## Supplementary material

**Supplementary File 1** Supplementary figures are attached after the main manuscript.

## Data availability

**Supplementary Data 1** A zip file containing the simulation executable, the source code, R scripts, batch files, and simulation results.

## SUPPLEMENTARY FIGURES

**Supplementary Figure 1.**
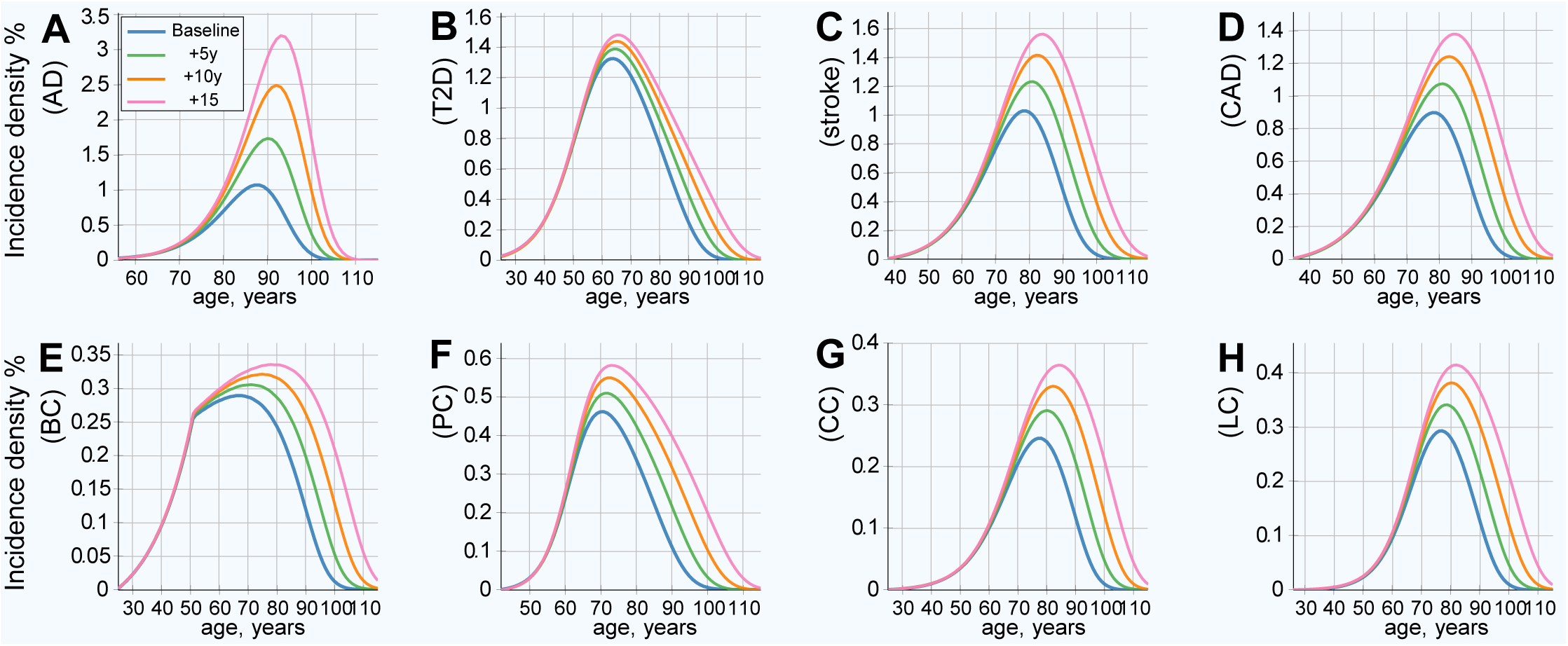
Baseline (without gene therapy) population cumulative incidence rate density. **(A)** Alzheimer’s disease, **(B)** type 2 diabetes, **(C)** cerebral stroke, **(D)** coronary artery disease, **(E)** breast cancer, **(F)** prostate cancer, **(G)** colorectal cancer, **(H)** lung cancer. The area under the curve is equal to the lifetime risk for each scenario, accounting for mortality. Projected LOD incidence rate relative to the number of individuals at birth, baseline, and scenarios with life expectancy increased by 5, 10, and 15 years. This is the baseline scenario without gene therapy or other health improvements for the plotted LOD. This represents the case where life expectancy increases due to causes other than the plotted LOD.

**Supplementary Figure 2.**
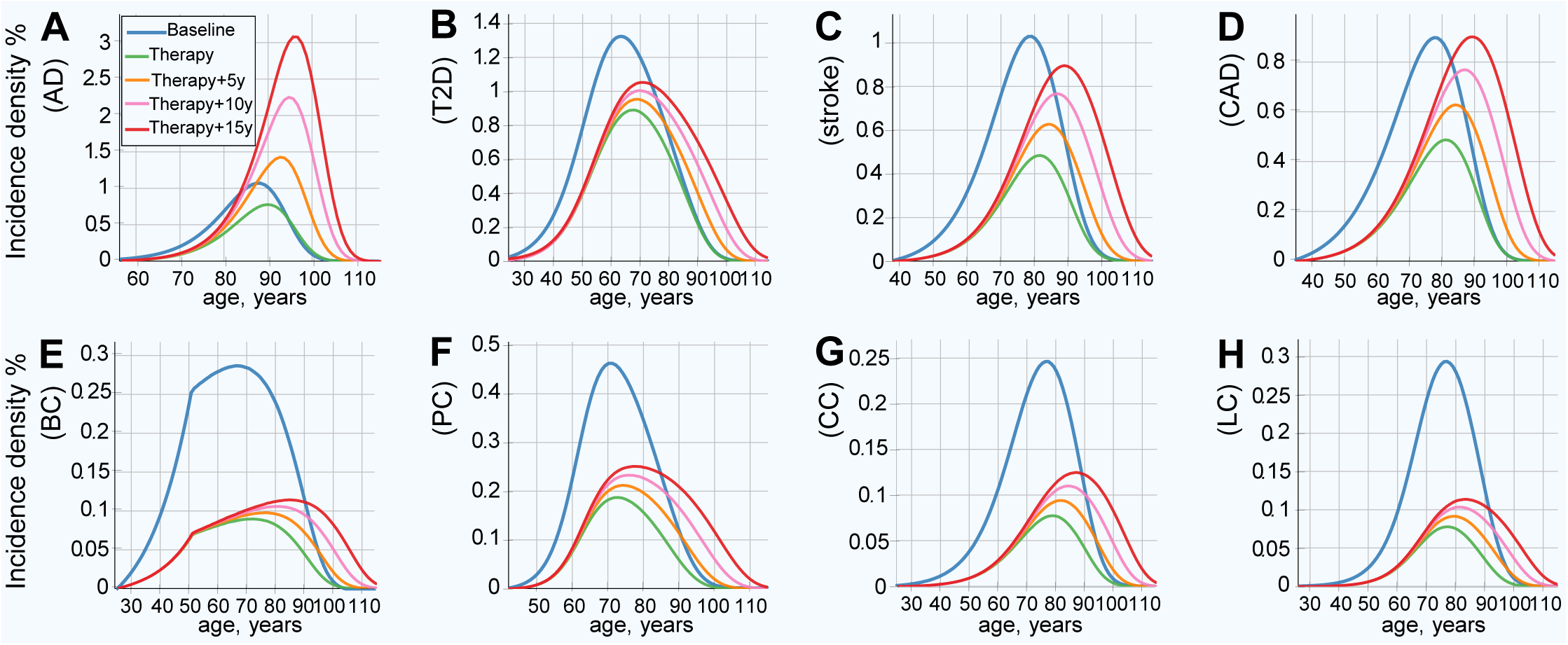
Population cumulative incidence rate density after emulated gene therapy. **(A)** Alzheimer’s disease, **(B)** type 2 diabetes, **(C)** cerebral stroke, **(D)** coronary artery disease, **(E)** breast cancer, **(F)** prostate cancer, **(G)** colorectal cancer, **(H)** lung cancer. All individuals in the population had emulated corrective gene therapy editing, on average, 15 SNPs (corresponding to an OR multiplier of 0.25). The area under the curve is equal to the lifetime risk for each scenario, accounting for mortality. Projected LOD incidence rate relative to the number of individuals at birth, after gene therapy, and scenarios with life expectancy increased by 5, 10, and 15 years.

**Supplementary Figure 3.**
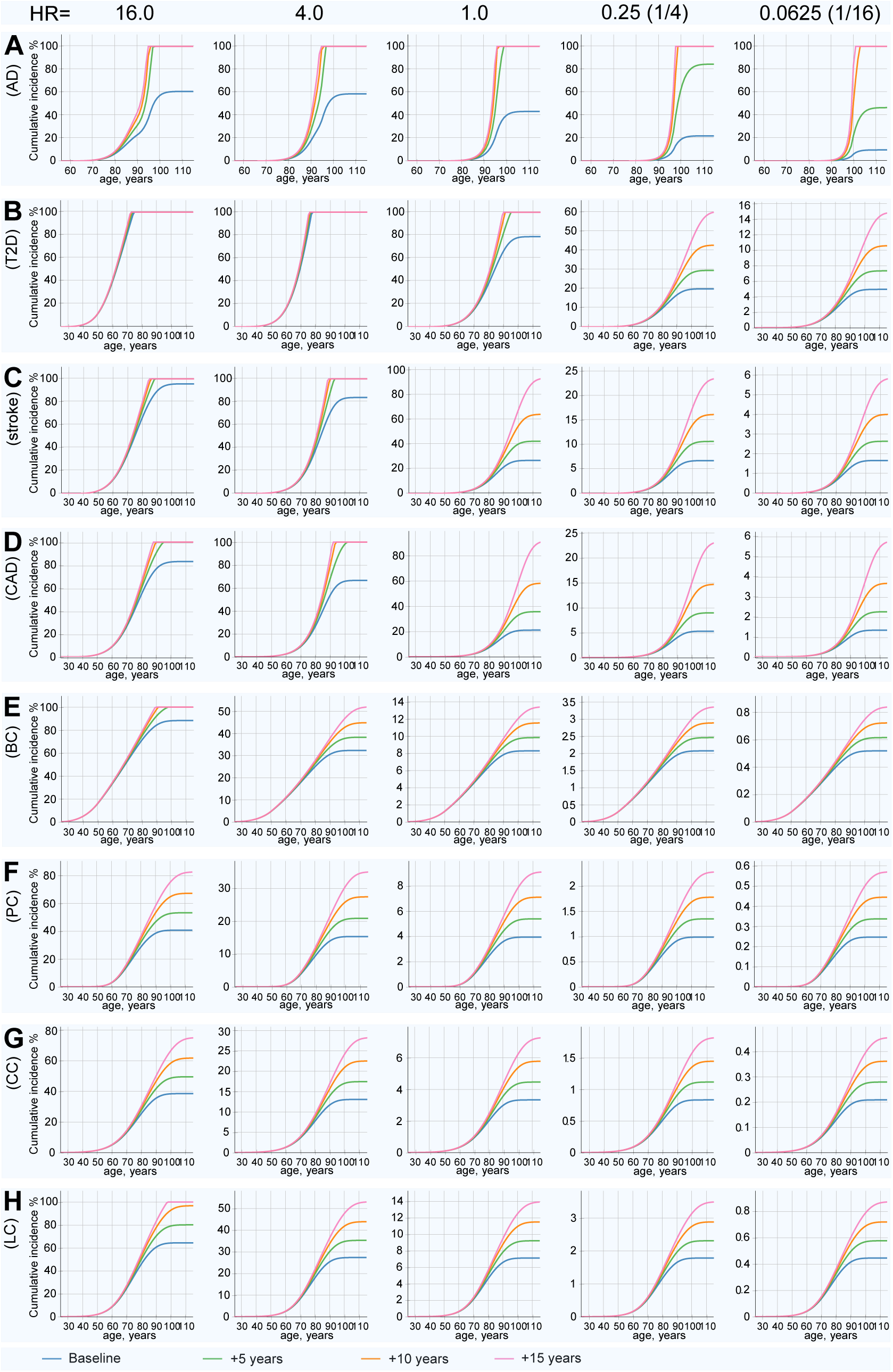
Matrix display of cumulative incidence and lifetime risk for discrete hazard ratio values. **(A)** Alzheimer’s disease, **(B)** type 2 diabetes, **(C)** cerebral stroke, **(D)** coronary artery disease, **(E)** breast cancer, **(F)** prostate cancer, **(G)** colorectal cancer, **(H)** lung cancer. This figure shows the matrix of calculated cumulative incidence for discrete hazard ratio (HR) values relative to the population mean, and cumulative incidence increase in 5-year life expectancy increments. Lifetime risk (lifetime cumulative incidence) corresponds to the lifetime (rightmost) values of the plots. For lower HRs, the lifetime values equal to the lifetime risk are almost precisely proportionate to the HR multiple (note the scale change), while the age progression curve for each HR value shifts toward older ages. This shift is most prominent for the highest incidence LODs; the change in scale moving from higher to lower HRs visually masks the shift appearance.

**Supplementary Figure 4.**
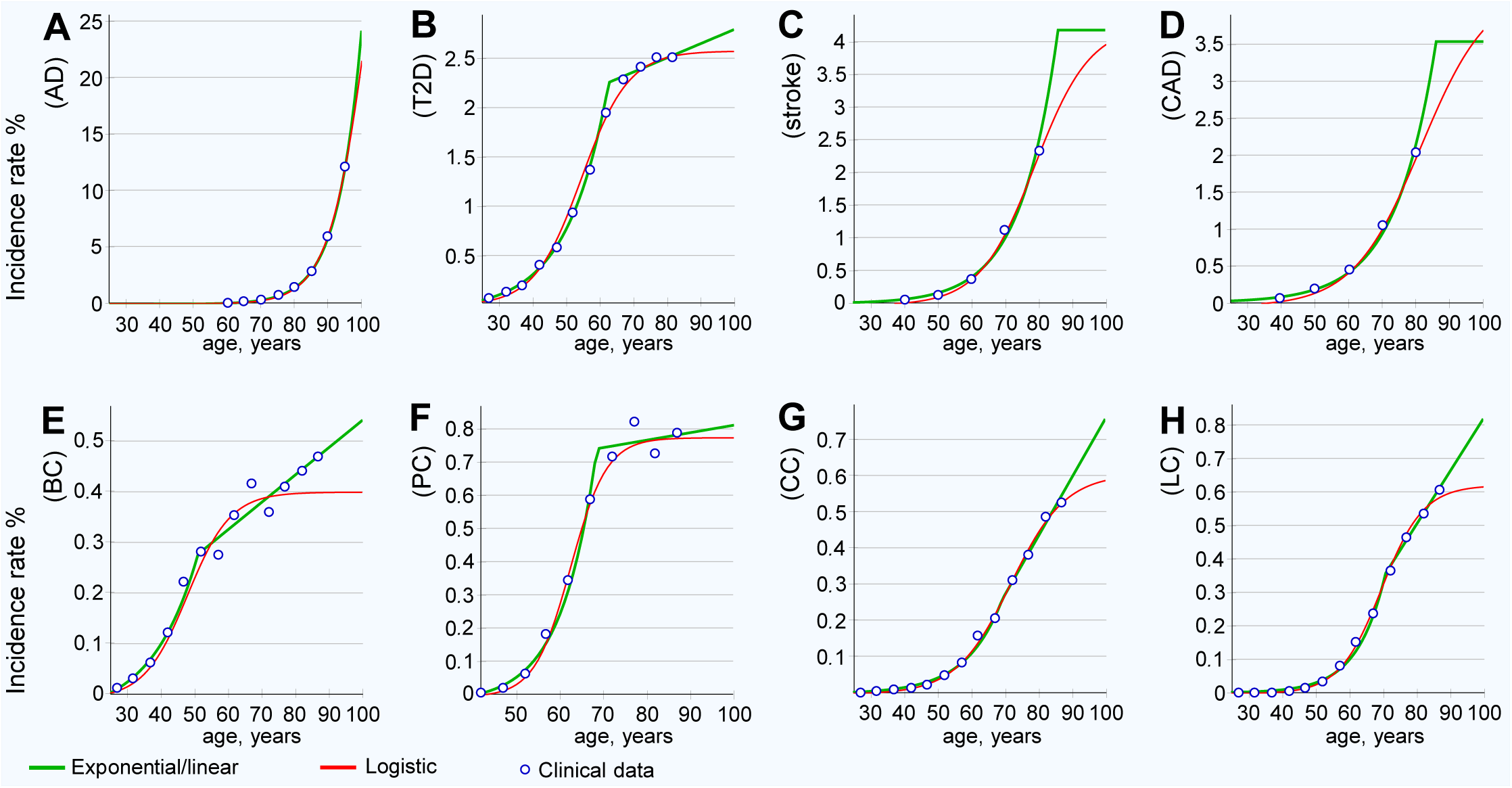
LOD clinical incidence rates and functional approximations. **(A)** Alzheimer’s disease, **(B)** type 2 diabetes, **(C)** cerebral stroke, **(D)** coronary artery disease, **(E)** breast cancer, **(F)** prostate cancer, **(G)** colorectal cancer, **(H)** lung cancer. Two functional approximations of clinical data: exponential followed by linear and logistic.

**Supplementary Figure 5.**
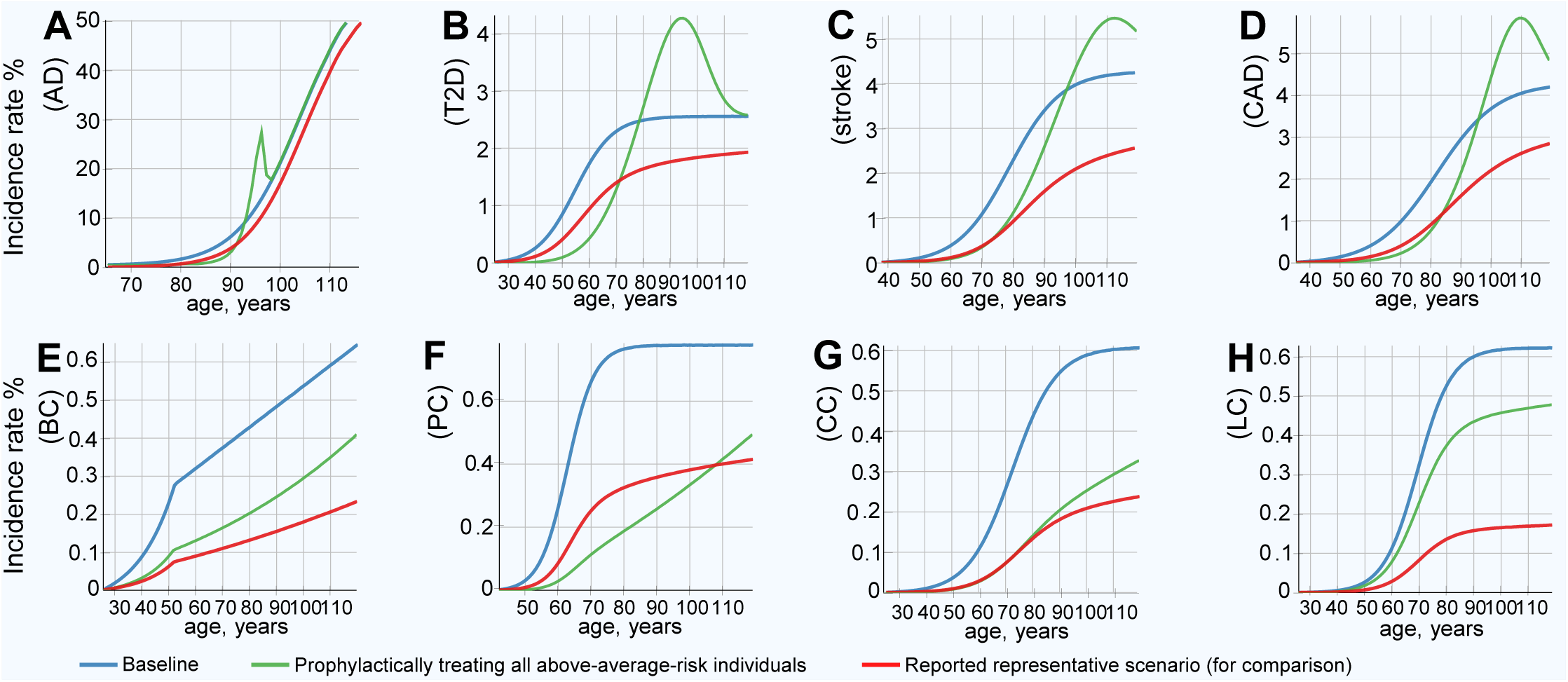
Incidence rate pattern, baseline and after gene therapy. Prophylactically treating above-average-risk individuals to match the polygenic risk average of the baseline population. **(A)** Alzheimer’s disease, **(B)** type 2 diabetes, **(C)** cerebral stroke, **(D)** coronary artery disease, **(E)** breast cancer, **(F)** prostate cancer, **(G)** colorectal cancer, **(H)** lung cancer. There is a pronounced delay in early incidence for all LODs. As a result of half of the population having PRSs equal to what was previously the population mean PRS, statistically, a large number of individuals are likely to become sick at relatively similar advanced age. For the four non-cancer LODs, there is a spike in the incidence rate at a very old age, which exceeds the baseline incidence rate. The spike appears steep; the proportion of the population that remains alive at this old age is diminishing. LODs with lower incidence and heritability do not display a spike exceeding the baseline incidence. The values reported in the article’s representative scenario are shown for comparison.

**Supplementary Figure 6.**
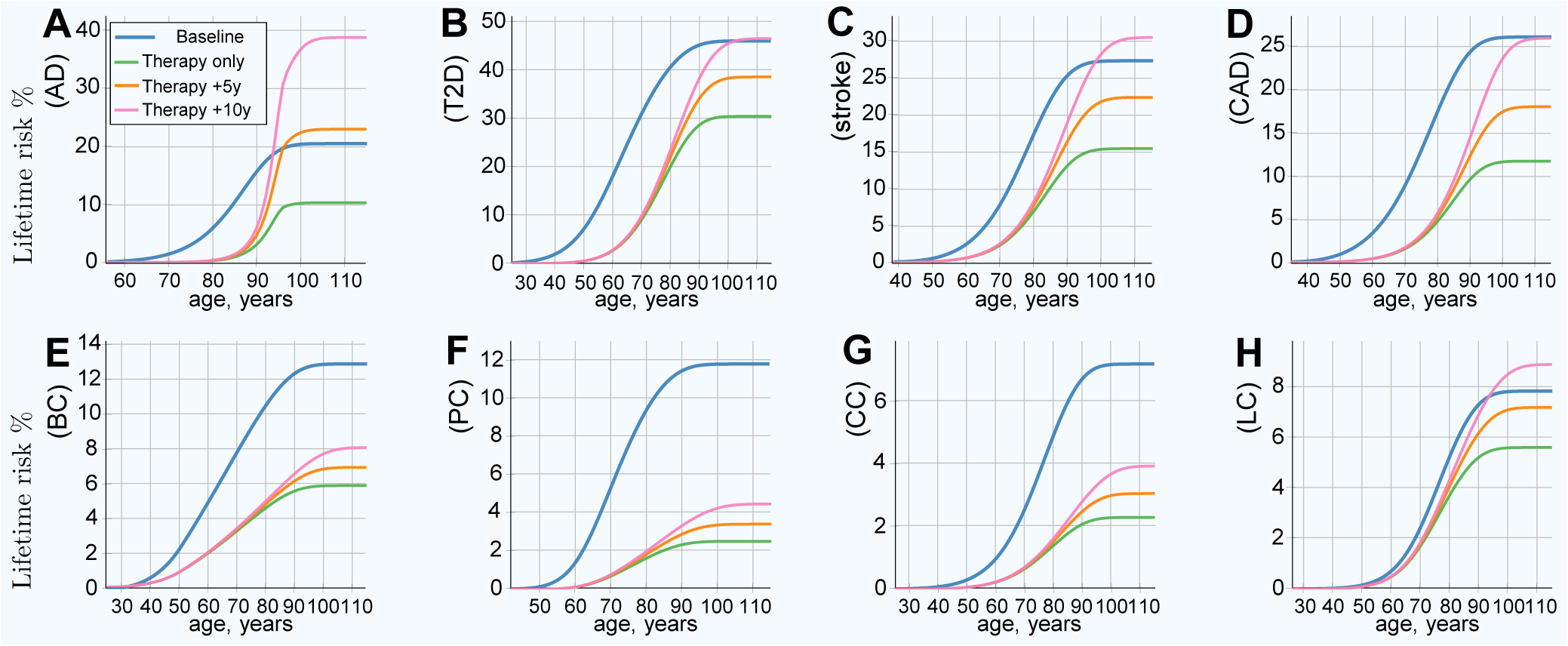
Lifetime risk, baseline and after gene therapy. Prophylactically treating above-average-risk individuals to match the polygenic risk average of the baseline population. **(A)** Alzheimer’s disease, **(B)** type 2 diabetes, **(C)** cerebral stroke, **(D)** coronary artery disease, **(E)** breast cancer, **(F)** prostate cancer, **(G)** colorectal cancer, **(H)** lung cancer. Showing lifetime risk after therapy, and the trends with life expectancy increased by 5 and 10 years. The lifetime risk results are qualitatively similar to the reported scenario, though not as large, and a life expectancy increase of 10 years results in the risk exceeding the baseline for the four non-cancer LODs and lung cancer. Lung cancer showed the least improvement. This is due to the disease having the lowest reported heritability, which results in the lowest PRS variance and the fewest edits for the treated population under this scenario.

